# UK mosquitoes are competent to transmit Usutu virus at native temperatures

**DOI:** 10.1101/2024.07.19.604126

**Authors:** Jack Pilgrim, Soeren Metelmann, Emma Widlake, Nicola Seechurn, Alexander Vaux, Karen L Mansfield, Jola Tanianis-Hughes, Ken Sherlock, Nicholas Johnson, Jolyon Medlock, Matthew Baylis, Marcus SC Blagrove

## Abstract

Usutu virus (USUV) is an emerging zoonotic virus transmitted primarily by *Culex* mosquitoes. Since its introduction into Europe from Africa during the late 20^th^ century, it has caused mortality within populations of passerine birds and captive owls, and can on occasion lead to disease in humans. USUV was first detected in the UK in 2020 and has become endemic, having been detected in either birds and/or mosquitoes every subsequent year. Importantly, the vector competence of indigenous mosquitoes for the circulating UK (London) USUV strain at representative regional temperatures is still to be elucidated. This study assessed the vector competence of five field-caught mosquito species/biotypes, *Culex pipiens* biotype *molestus*, *Culex pipiens* biotype *pipiens*, *Culex torrentium*, *Culiseta annulata* and *Aedes detritus* for the London USUV strain, with infection rates (IR) and transmission rates (TR) evaluated between 7 to 28 days post-infection. Infection and transmission were observed in all species/biotypes aside from *Ae. detritus* and *Cx. torrentium*. For *Cx. pipiens* biotype *molestus*, transmission potential suggests these populations should be monitored further for their role in transmission to humans. Furthermore, both *Cx. pipiens* biotype *pipiens* and *Cs. annulata* were shown to be competent vectors at 19°C indicating the potential for geographical spread of the virus to other UK regions.

## Introduction

Over the last century, the United Kingdom has not been severely impacted by mosquito-borne diseases, with the last outbreak of locally transmitted malaria occurring in the early 1920s [1]. However, in 2020 the first recorded case of Usutu virus (USUV) was confirmed in five dead Eurasian blackbirds (*Turdus merula*) and one house sparrow (*Passer domesticus*) from London Zoo [2]. Usutu virus is a species of *Orthoflavivirus* which spread to Europe from Africa during the late 20^th^ century [3,4] and is thought to be transmitted primarily by *Culex* spp. [5]. Similar to West Nile Virus (WNV), the primary reservoir for USUV is birds, with the orders Passeriformes and Strigiformes over-represented in displaying clinical signs [6]. However, spillover to mammals, including humans, has been reported and can occasionally cause neurological symptoms [7]. Since its initial detection in the UK, USUV has been detected annually through both bird and mosquito surveillance [8–11]. Usutu virus is now endemic in the UK, as inferred from molecular clock analysis of index cases and subsequent cases, indicating it has successfully overwintered [10]. The full extent of this emerging threat to the UK is still under consideration, although there is circumstantial evidence suggesting USUV is impacting wild bird populations, with a decline in reported sightings of blackbirds in London corresponding with the initial outbreak [11]. A report from 2023 has suggested the geographic range of the virus has expanded to include parts of Cambridgeshire north of the index site [9]. Therefore, a greater understanding of indigenous mosquito competence and the thermal limits of the virus is of pressing need to understand the future impact of USUV in the UK.

The main implicated vector of USUV in Europe is the *Culex pipiens* complex, which includes the morphologically indistinguishable forms *Culex pipiens* biotype *pipiens* and *Culex pipiens* biotype *molestus* (hereafter *Cx. pipiens* and *Cx. molestus*) [12]. *Culex molestus* exhibits several behavioural differences to *Cx. pipiens* with the former being autogenous (able to lay eggs without a blood-meal [13]), stenogamous (able to mate in confined spaces [14]) and mammalophilic [15]. In comparison, *Cx. pipiens* mates in open spaces, generally feeds on birds and mated females require a bloodmeal in order to lay their first egg batch [16]. These distinct behavioural characteristics implicate both biotypes as having unique putative roles to play in transmission dynamics of USUV. *Culex molestus* are active all year around and generally live in underground human-made structures such as flooded basements and sewage treatment works where they may be perceived as a biting nuisance [17]. However, the emergence of USUV now necessitates that *Cx. molestus* be investigated for their potential to transmit disease to the public. In contrast, *Cx. pipiens* is likely to contribute to virus amplification and death in birds through its ornithophilic feeding patterns. There has also been a suggestion that the ability of both *Cx. pipiens* and *Cx. molestus* to hybridize increases the risk of spillover due to the production of vectors with more generalized feeding on both birds and humans (referred to as “bridge vectors”) [18–20].

Vector competence, the ability of a vector to transmit an infectious agent after exposure, is impacted by a range of factors. These include (but are not limited to) the strain and titre of the virus [21–23], genetic variability of the insect host [21,24], and temperature [25], which influences the extrinsic incubation period of the virus. There have been several USUV vector competence studies conducted in Europe which have found *Culex modestus*, *Cx. pipiens*, and *Cx. molestus* capable of laboratory transmission of the virus [26–30]. However, to date, the only study investigating the role of indigenous UK mosquitoes found very low USUV transmission potential [31]. This was observed in two colonies—one *Cx. pipiens* and one hybrid—incubated at 25°C using the SAAR 1776 strain (originating from South Africa). Only a single specimen from the hybrid colony tested positive for USUV in its saliva after virus challenge. Testing a distantly related virus strain at high temperatures might not account for the dynamics of the UK endemic strain, which operates at lower temperatures compared to many USUV transmission areas in mainland Europe. In light of this, the current study assessed the ability of five UK mosquito species/biotypes to transmit the London 2020 USUV strain under current UK climatic conditions. These were *Cx. molestus*, *Cx. pipiens* and *Cx. torrentium*, alongside other potential bridge vectors *Aedes detritus* and *Culiseta annulata* [32].

## Methods

### Mosquito collections

Five species/biotypes of mosquitoes were collected from Cheshire and Greater London regions of the UK: *Culex pipiens pipiens, Culex pipiens molestus, Culex torrentium, Culiseta annulata and Aedes detritus* (Figure 1). The morphologically indistinguishable biotypes, *Cx. pipiens, Cx. molestus* and *Cx. torrentium,* along with *Cs. annulata* were collected as egg rafts. Approximately 200 *Cx. molestus* egg rafts were collected from a large municipal Wastewater Treatment Facility located in West London on the 1^st^ August 2022 and transferred to the climate-controlled insectary (Leahurst Campus, University of Liverpool, Neston, UK) and maintained at 24°C, 12-h light/12-h dark and 80% Relative Humidity (RH). Six *Cx. molestus* egg rafts were allowed to hatch per 35 x 25 x 5 cm larval tray with 2 litres of water. For nutrition, 125 mg Brewer’s Yeast (Holland’s and Barrett, UK) and 500 mg Koi carp pellets (Extra Select, UK) were added every 2 days until signs of pupation. *Culex pipiens, Cx. torrentium* and *Cs. annulata* were collected from container habitats from Ness Botanic Gardens, Neston, Cheshire (53°16’25.1724“N, 3°2’42.2736”W) and a nearby farm at the University of Liverpool, Leahurst Campus, Neston, Cheshire (53°17′5.3″N, 3°01′33.3″W) between June-September 2023. Hatched larvae of *Cx. pipiens* and *Cs. annulata* were reared in the same manner as *Cx. molestus*, except they were housed in a non-insulated brick outbuilding to mimic outdoor shaded conditions during the UK vector season. *Aedes detritus* were collected as fourth-instar larvae or pupae from salt marshes in Little Neston, Cheshire (53°16′37.2″N, 3°04′06.4″W) between August and October 2022. Adults were allowed to emerge in 30 × 30 × 30 cm BugDorm cages (BugDorm, Taichung, Taiwan) and offered 10% sucrose solution on cotton wool alongside *ad libitum* water. For later infection work, *Culex pipiens* biotypes (*Cx. molestus* and *Cx. pipiens*) were provisionally allocated based on ecological niche characteristics (underground vs overground habitats) before confirmation using a molecular assay. Specifically, an initial PCR was used to distinguish between *Cx. torrentium* and *Cx. pipiens*/*Cx. molestus* using a restriction enzyme assay targeting the 3’ end of the COI gene [33]. Subsequently, a multiplex PCR was employed to differentiate between *Cx. pipiens*, *Cx. molestus* and hybrids [34].

**Figure 1.**
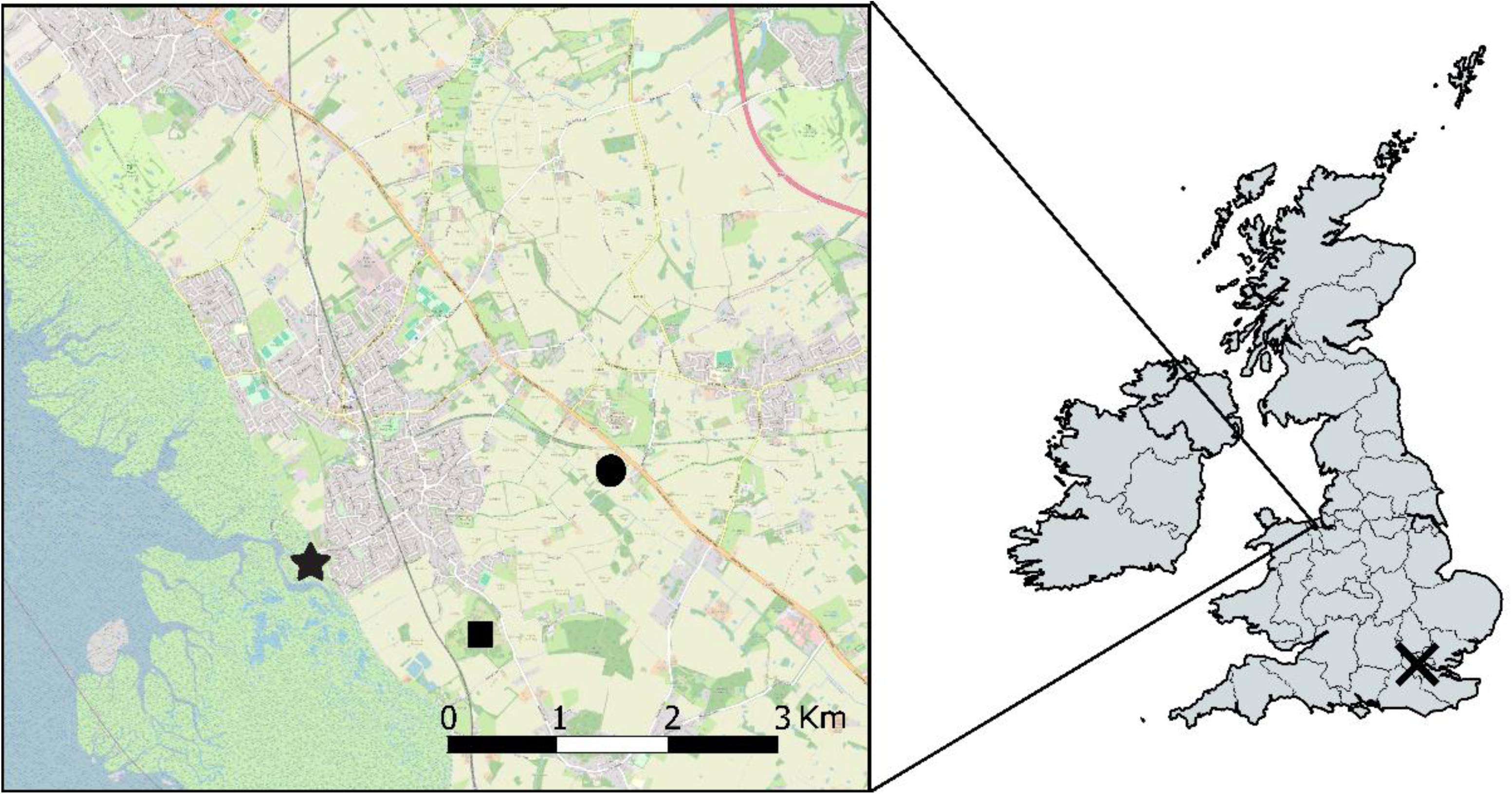
Map showing the different mosquito collections sites in Cheshire (inset) and London between 2022 and 2023. Star = *Aedes detritus*, Square = *Culiseta annulata and Culex pipiens pipiens,* Circle = *Culex pipiens pipiens,* Cross (London) = *Culex pipiens molestus*. The map was generated with QGIS version 3.36.2 [35] using OSM standard as the base map [https://wiki.openstreetmap.org/]

### Virus

USUV strain London 2020 was originally isolated from the brain and kidney of an infected black bird in London (GenBank accession number: MW001216) [2], with isolated virus propagated through four passages in Vero cells. The titre of stock virus of kidney origin was confirmed by plaque assay in Vero cells as 4.0 x 10^8^ PFU/ml. Virus stocks maintained at the APHA (The Animal and Plant Health Agency, Surrey, UK) were transported to the University of Liverpool, Leahurst campus and aliquoted on the day of receipt before storage at −80°C.

### Infection experiments

Twenty-four hours before viraemic blood feeding, access to sugar and water was removed. At 7-14 days post-emergence, adult mosquitoes were transferred into smaller BugDorm cages (17.5 x 17.5 x 17.5 cm). They were then fed for four hours in low light conditions at 24°C on defibrinated horse blood (Scientific Laboratory Supplies, Nottingham, UK) containing 2 mM ATP and a final titre of 1 × 10^7^ PFU/mL of virus through dilution of aliquot stocks. To achieve this, a Hemotek feeder (Discovery *Workshops*, Lancashire, UK) heated to 39 °C was used with chick skins as membranes and bird feathers collected from the external environment to stimulate probing behaviour. Blood-fed females were incubated at 17°C, 19°C, 21°C, or 23°C with a relative humidity (RH) of >90% and a 12-hour light/12-hour dark cycle. These temperatures were selected based on Met Office data [36], with the highest UK daily average maximum temperature recorded between 1991 and 2020 being 23°C during the months of July and August. Mosquitoes were maintained at these temperatures for 7-28 days depending on the numbers available for feeding and were provided with 10% sucrose which was changed every two days.

Mosquitoes were then anaesthetised with FlyNap (triethylamine; FlyNap, Carolina Biological Supply Company, Burlington, USA), before saliva expectorate was collected by inserting each mosquito’s proboscis into a capillary tube containing mineral oil for 30 minutes. Each mosquito was checked for salivation under a microscope through observation of an air bubble at the tip of the labium. Expectorates and bodies were then placed in 500 µl TRIzol^TM^ reagent (Fisher Scientific, Loughborough, UK), before storage at −80°C. Zero day post-infection (dpi) mosquitoes were used as positive controls.

### Measuring viral RNA in bodies and saliva

For both bodies and saliva, RNA was extracted using TRIzol^TM^ according to the manufacturer’s instructions. The DNA interphase of the phenol:chloroform mixture was retained for *Culex pipiens* biotype identification as described above. Quantitative RT-PCR was then used to quantify viral RNA genome copies in both mosquito saliva and bodies. To this end, the Qiagen Quantitect probe RT-PCR kit (Qiagen, Manchester, UK) was used according to the manufacturer’s instructions. TaqMan quantitative polymerase chain reaction (qPCR) was used alongside NS1 primers which were newly designed to complement the London 2020 strain: USUTU_Lon_3177 F (5’-CGTGAAGGTTACAAAGTCCAGA; nucleotide positions 3177–3198 for GenBank accession no. MW001216) and USUTU_Lon_3284 (5′-TCTTATGGAGGGTCCTCTCTTC; nucleotide positions 3284–3305 for GenBank accession no. MW001216). A TaqMan probe (Fisher Scientific, Loughborough, UK) and 6-FAM reporter dye were used along with a QSY quencher dye with the probe sequence as described in Jost et al. [37]. Samples were run in duplicate on a Roche LightCycler 480 with the mean of the two C_q_ values being quantified using absolute quantification. PCR plates were incubated at 50°C for 30 min, then 95°C for 15 min (reverse transcriptase step) before cycling for 45 cycles at 94°C for 15 seconds followed by the annealing/extension step at 60°C for 1 min.

### Analytical sensitivity of assay

Given the typically low titre levels of USUV in expectorate [29], only a single qRT-PCR assay was chosen to maintain sensitivity, rather than splitting samples between both plaque and molecular assays [38]. In addition, a prior study has demonstrated a significant correlation between qRT-PCR and plaque assay techniques for Flaviviruses [39]. To assess the sensitivity of the assay, a standard curve was generated using synthetic oligonucleotides provided by Integrated DNA Technologies (IDT, Solihull, UK) through their Gblocks service, which supplied quantified copy numbers of the 128bp amplicon produced by the PCR. Standards were serially diluted by a factor of 10, ranging from 1.7 x 10^12^ copies/µl to 1.7 x 10^1^ copies/µl. The minimum detection limit was determined to be a cycle threshold (Ct) of 38.1 at 1.7 x 10^1^ copies/µl. Consequently, detection at a maximum of 38 cycles was set for a positive result.

### Data analysis

The percentage of bodies with detectable viral RNA determined infection rate (IR), whereas the percentage of saliva positives from bodies determined transmission rate (TR) at any given time point. All statistical analyses were performed in Rstudio V2022.02.2 [40]. The difference in IR and TR proportions at each time point and temperature was tested for statistical significance using Fisher’s Exact test. For significant differences in viral titres between species at the same time point and temperature, the Wilcoxon-Rank Sum test was used. For comparing titres between time points for the same species and temperature, the Kruskal-Wallis test was utilised. Statistical significance was determined using a two-tailed *p*-value threshold of 0.05

### Temperature suitability mapping

Finally, gridded climate data from the HadUK-Grid dataset [41] for the period between 2018 and 2022 was used to investigate areas suitable for USUV transmission by *Cx. pipiens*. Based on empirical data from the vector competence experiments, locations where average temperatures over a 14-day period reached or exceeded the minimum threshold for USUV transmission were identified, and days with possible transmission risk (TRD) calculated. These areas were then mapped using a 5 km grid resolution to highlight potential transmission zones and to validate that the vector competence-temperature data were consistent with field detections of USUV in the UK (London Zoo, Greater London, and Cambridgeshire; unknown site with the centre of Cambridgeshire used for mapping). The resulting TRD were averaged over the five-year period to identify consistent patterns, with zones meeting the transmission criteria annually also marked.

## Results

### UK field populations of *Culex pipiens pipiens* and *Culex pipiens molestus* demonstrate USUV transmission potential

Both *Culex pipiens* biotypes (*Cx. molestus* and *Cx. pipiens*) were used to investigate the vector competence of field-caught populations for the London strain of USUV. Molecular species identification confirmed the virus challenge and end point survival of 237 *Cx. pipiens* from the Cheshire region and 189 *Cx. molestus* from the Greater London region. Additionally, a limited number of *Cx. pipiens*/*molestus* hybrids (N=4, Cheshire; N=7, London) and *Cx. torrentium* (N=5, Cheshire) were used for infections (Table 1). Five *Cx. molestus* were found in the Cheshire catches and removed from analysis (Supplementary Table S1).

**Table 1.**
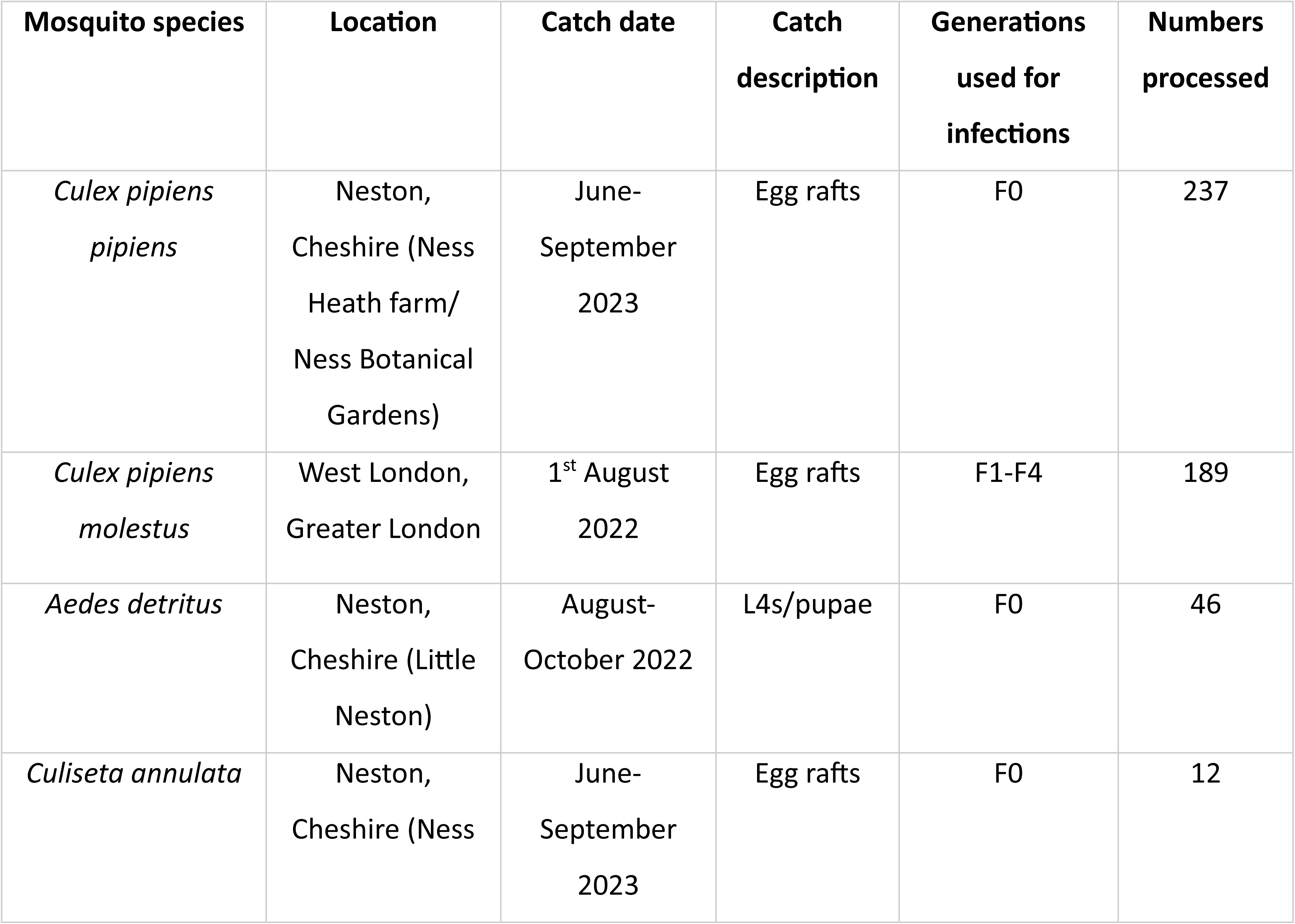
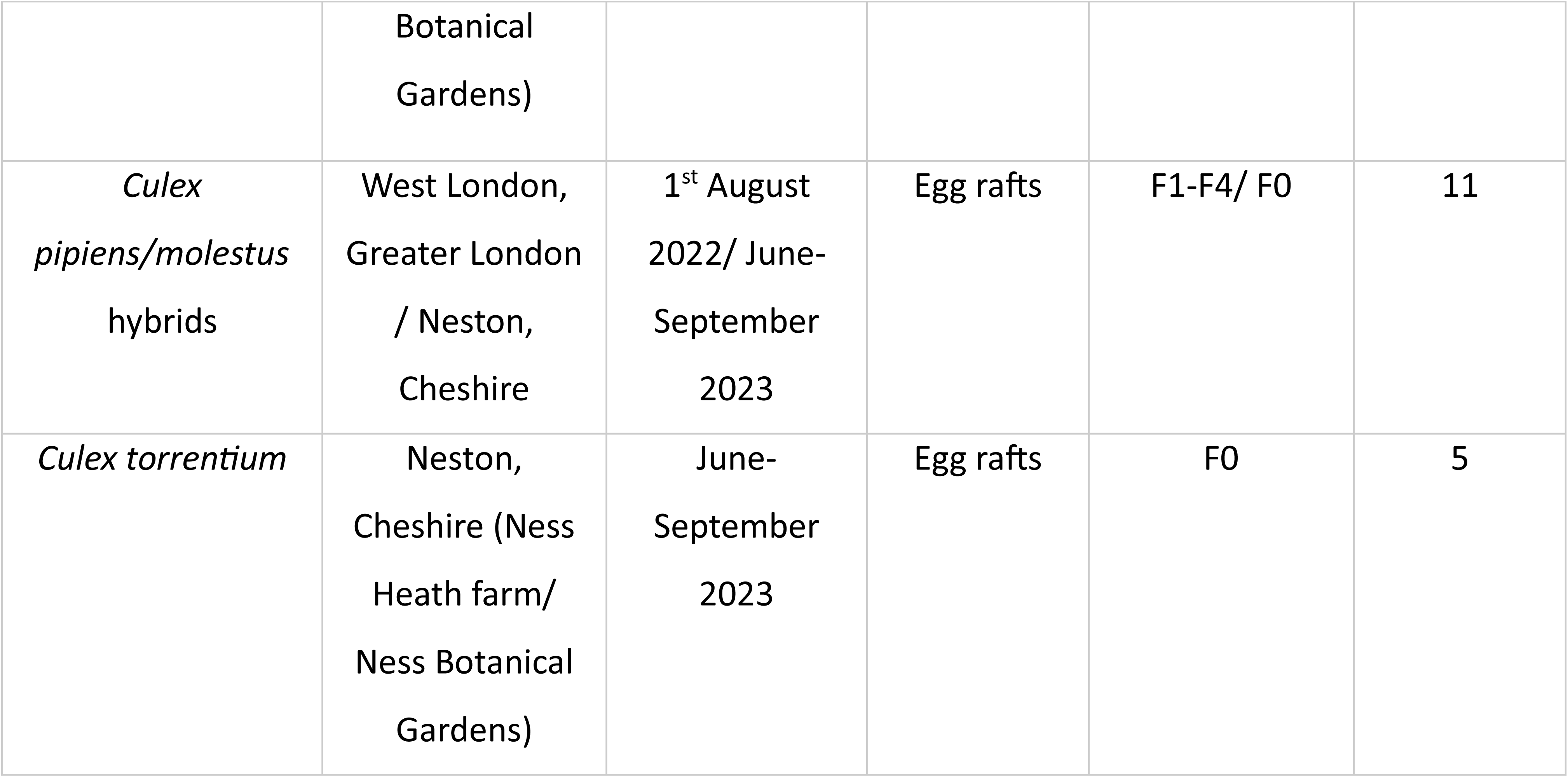
Collection details of mosquito species used for USUV infection work.

Virus was detected in mosquito bodies at all temperatures for *Cx. pipiens* and *Cx. molestus* except 23°C at 28 days post-infection (dpi) for *Cx. pipiens* (Figure 2C). For *Cx. molestus*, a similar pattern was seen for all temperatures during infection experiments. Early infection rates at seven dpi ranged from 20-38%, before troughing at 14 dpi (IR range=9-21%), and then recovering to 27%-91% at 28 dpi. Despite this, the only significant differences observed between time points within the same temperature were at 23°C (Figure 2A). Specifically, the IR at 21 dpi (75.0% IR, *N* = 20) was significantly higher compared to 7 dpi (20.0% IR, *N* = 20, Fisher’s Exact: p = 0.0012) and 14 dpi (15.8% IR, *N* = 19, Fisher’s Exact: *p* = 0.0003). Similarly, the IR at 28 dpi (90.9% IR, *N* = 11) was significantly higher compared to 7 dpi (20.0% IR, *N* = 20, Fisher’s Exact: *p* = 0.0003) and 14 dpi (15.8% IR, *N* = 19, Fisher’s Exact: *p* = 0.00009).

**Figure 2.**
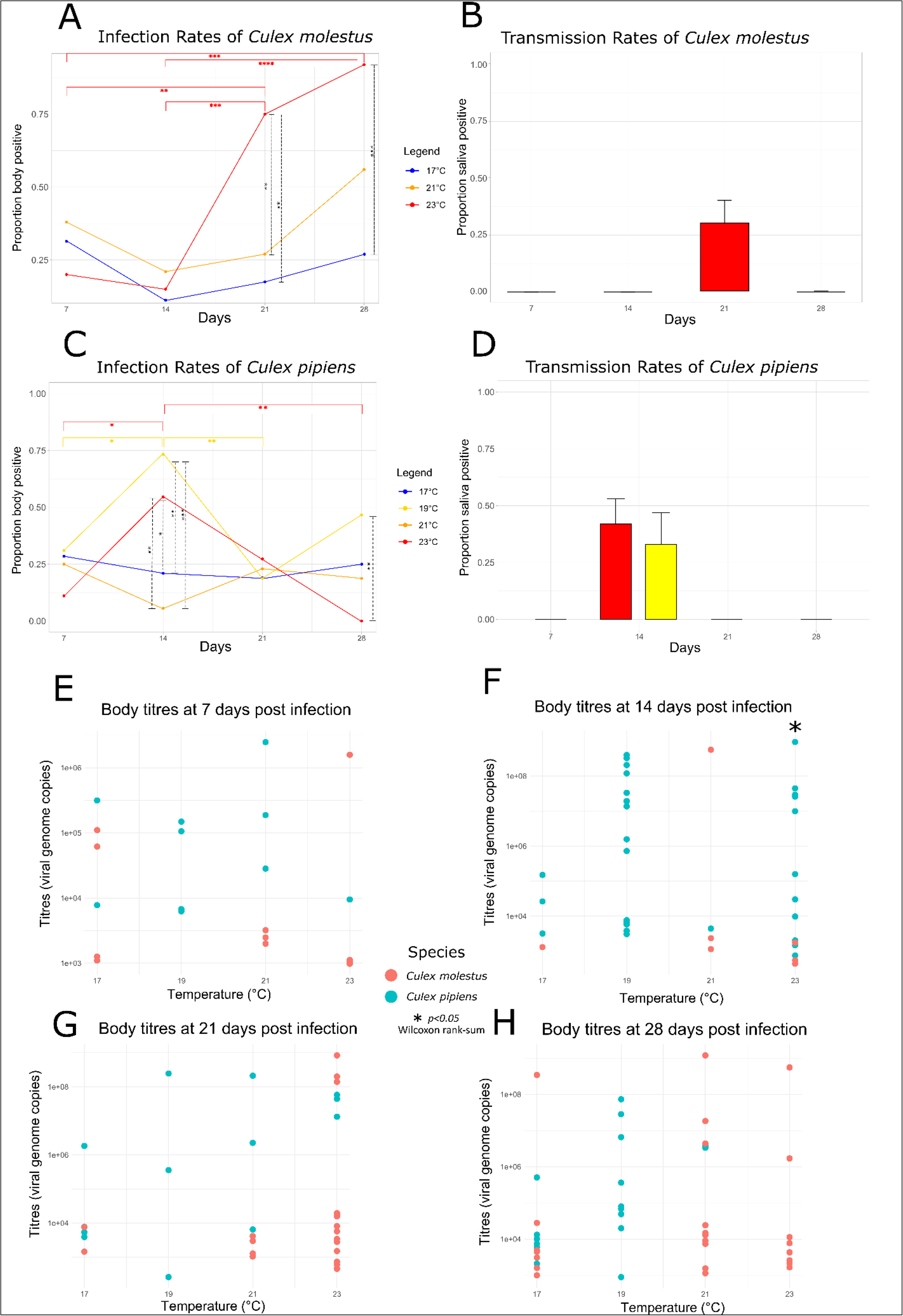
The proportion of total *Cx. molestus* and *Cx. pipiens* bodies (A,C) and saliva (B,D) positive for USUV at four temperatures (17°C, 19°C, 21°C, 23°C) and four time points (7, 14, 21 and 28 days post-infection). The difference in IR and TR proportions at each time point and temperature was tested for statistical significance using Fisher’s Exact test with *p*-value thresholds:* < 0.05, ** < 0.01, *** < 0.001, **** < 0.0001. USUV body titres of *Cx. molestus* and *Cx. pipiens* at 7 (E), 14 (F), 21 (G) and 28 (H) dpi were tested for significant differences using the Wilcoxon rank-sum test with a p-value threshold of 0.05. Numbers of mosquito individuals at each temperature and time point is shown in Supplementary Table S1.

Furthermore, these later time points at the highest temperature of 23°C gave significantly greater proportions of infected mosquitoes compared to the two cooler temperatures of 17°C and 21°C. Specifically, at 23°C, the infection rates at 21 dpi (75.0%, *N* = 20) were significantly higher than at 17°C (16.7%, *N* = 12, Fisher’s Exact *p* = 0.003) and 21°C (26.7%, *N* = 15, Fisher’s Exact *p* = 0.007). Similarly, at 23°C, the infection rates at 28 dpi (90.9%, *N* = 11) were significantly higher than at 17°C (26.9%, *N* = 26, Fisher’s Exact *p* = 0.0007) and 21°C (55.6%, *N* = 18, Fisher’s Exact p = 0.02). Despite the increasing infection rate over time, only the 21 dpi timepoint at 23°C produced saliva positives (30.0%, *N* = 20) with all other temperatures and timepoints failing to demonstrate transmission potential (Figure 2B).

Compared to *Cx. molestus*, no discernible pattern across temperatures was observed for *Cx. pipiens* infections. The highest proportion of infected bodies was observed at 14 dpi for 19°C (73.6%, *N* = 19) and 23°C (55.0%, *N* = 20) interventions. At 14 dpi the IRs at 19°C was significantly higher compared to the same time point at 17°C (21.4%, *N* = 14, Fisher’s Exact: *p* = 0.005) and 21°C (6.7%, *N* = 15, Fisher’s Exact: *p* = 0.0001). Similarly, the IR at 14 dpi for 23°C was significantly higher compared to the same time point at 17°C (21.4%, *N* = 14, Fisher’s Exact: *p* = 0.046) and 21°C (6.7%, *N* = 15, Fisher’s Exact: *p* = 0.004). By comparison, 21°C and 17°C interventions remained at a near constant low/intermediate IR (6.7-28.5%) throughout the experiment. In contrast to *Cx. molestus* which reached 90.1% IR (*N* = 11), *Cx. pipiens* infection rates at 23°C dropped to 0% at 28 dpi (*N* = 12). Saliva positives were detected at 14 dpi (Figure 2D) at both 19°C (37.5%, *N* = 16) and 23°C (41.7%, *N* = 12), with no additional occurrences found at any other time points. For both *Cx. molestus* and *Cx. pipiens*, there was no significant difference between body or saliva titres at any time point or temperature, apart from bodies from both species at 23°C and 14 dpi (Wilcoxon Rank Sum: *p* = 0.022; Supplementary Table S1). Additionally, for both species there were no body titre differences between time points at each temperature intervention (Kruskal-Wallis *p* > 0.05; Supplementary Table S1). However, active virus replication determined by titres detected above the average titres for 0 dpi specimens, was detected at every temperature for both *Cx. pipiens* and *Cx. molestus* bodies (Supplementary Figure S1 A-G). Five out of eleven *Cx. pipiens*/*molestus* hybrids and 3/5 *Cx. torrentium* demonstrated body infections (Supplementary Table S1) but none produced saliva positives.

### *Culiseta annulata* are able to transmit USUV but *Aedes detritus* appear to be refractory to USUV infection

A total of 12 *Culiseta annulata* survived until the end of the infection experiments after being challenged with USUV. However, due to the low initial number of specimens challenged, observations were made only at a single time point of 14 dpi and under three temperatures (17°C, 21°C, and 23°C). At 21°C, two out of seven bodies were infected, while one individual also demonstrated infection at 6,400 viral copies in its expectorate (Supplementary Figure S1 H). Individuals at 17°C (*N* = 3) and 23°C (*N* = 2) showed no signs of infection or transmission. Forty-six *Aedes detritus* assessed at similar temperatures also demonstrated no signs of infection or transmission (Table 2).

**Table 2.** The infection and transmission rates of Usutu virus in two mosquito species, *Aedes detritus* and *Culiseta annulata*, at 14 days post infection. ND=No data.

| Species | Infection rates (% positive) |  |  |  | Transmission rates (% positive) |  |  |  |
| --- | --- | --- | --- | --- | --- | --- | --- | --- |
|  | 17°C | 19°C | 21°C | 23°C | 17°C | 19°C | 21°C | 23°C |
| *Aedes detritus* | ND | 0/17 | 0/13 | 0/16 | ND | 0/17 | 0/13 | 0/16 |
| *Culiseta annulata* | 0/3 | ND | 2/7<br>(28.6%) | 0/2 | 0/3 | ND | 1/7<br>(14.3%) | 0/2 |

### Temperature suitability for USUV transmission by *Culex pipiens pipiens* in the UK

To validate that the vector competence results were consistent with field detections of USUV in the UK, geographical areas identified as within the thermal limits for USUV transmission were mapped. This was done by focusing on *Cx. pipiens*, the most common mosquito in the country and associated with USUV outbreaks. Historical temperature data across the country were analysed to identify regions where the average temperature over a period of 14 consecutive days reached or exceeded 19°C. After the initial 14-day period, each subsequent day was considered a potential Transmission Risk Day (TRD). TRDs were calculated for each year from 2018 to 2022. The resulting map (Figure 3) highlights regions that correspond with known USUV outbreak areas. In addition, regions with perceived suitable temperature conditions for USUV transmission include parts of the North West of England, South Yorkshire and the Midlands.

**Figure 3.**
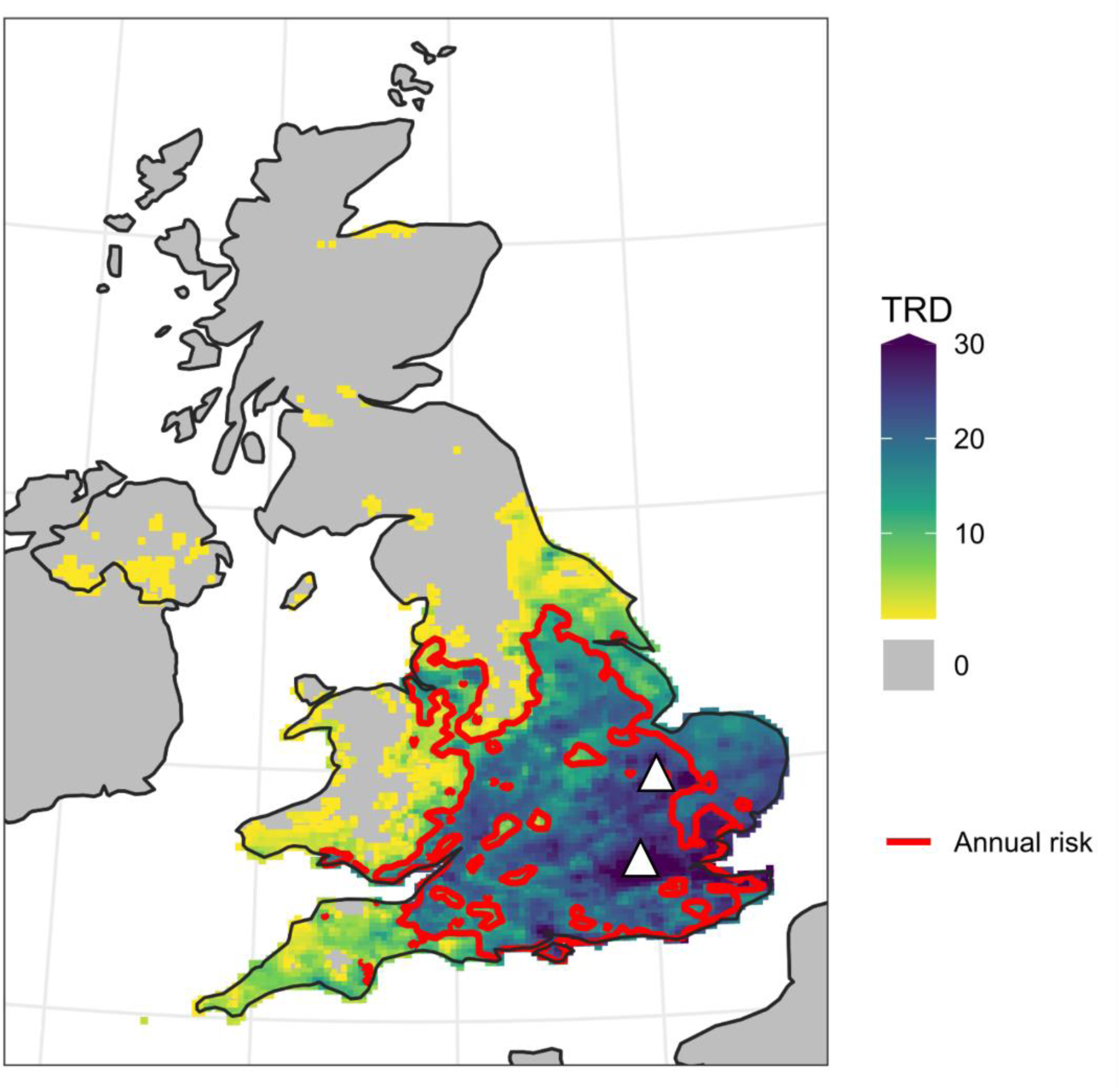
Transmission risk map of the UK from 2018 to 2022, highlighting regions within the thermal limits for *Cx. pipiens* Usutu virus (USUV) transmission. The coloured areas indicate the average number of days per year that reached or exceeded the minimum temperature threshold for USUV transmission (19°C) following a sufficiently warm 14-day period, referred to as Transmission Risk Days (TRDs). Regions meeting this criterion annually are outlined with a red contour line. White triangles=known USUV outbreaks. Ireland and France are not analysed rather than no risk.

## Discussion

This study is the first to assess the susceptibility and transmission of the endemic London strain of USUV in indigenous populations of UK mosquitoes. Due to differences in host preference and habitats, separating the *Culex pipiens* complex into its two biotypes, *molestus* and *pipiens*, is essential for understanding their role in USUV transmission dynamics. This separation also aids in determining the likely human exposure to circulating strains of USUV, with *Cx. pipiens* identified as the enzootic vector and *Cx. molestus* as a potential bridge vector. For *Cx. molestus*, the sharp increase in infection rates (75%) at 21 dpi and 23°C coinciding with the presence of the virus in saliva is noteworthy. This is comparable to a previous study by Holicki *et al*. [27] where similar rates of body infections were observed in two colonies of *Cx. molestus* (67% and 80%), alongside transmission at a similar temperature (25°C). In the same study, transmission was also observed between 14-16 dpi whereas no other time point aside from 21 dpi demonstrated transmission potential for our study. This discrepancy in infection and transmission dynamics witnessed in *Cx. molestus* is also observed in a recent study by Krambrich et al. [30] which, like Holicki *et al.* [27], also found transmission occurring at earlier times post infections (7 dpi and 14 dpi) but demonstrated low infection rates at 28 dpi (6%, *N* = 42) compared to our findings of 91% (*N* = 11). This inconsistency in findings for *Cx. molestus* epitomises the problem of extrapolating findings from vector systems non-representative of the target vector population, environment and virus of interest [42]. For example, the use of mosquitoes from different countries with genetic heterogeneity could explain such observed differences; Holicki *et al.* [27] used *Cx. molestus* colonies from Serbia and Germany, while Krambrich *et al*. [30] used a colony originating from Sweden. Furthermore, the utilization of Bologna/09 [30] and ME/2015/ED-A [27] strains in the aforementioned studies (which have 29 and 41 amino acid differences in their polyproteins compared to the London 2020 strain) could also explain these discrepancies.

The ability of *Cx. molestus* to transmit USUV is of interest due to their mammalophilic feeding behaviour, potentially implicating the species in the transmission of the virus to humans. [32]. *Cx. molestus* gained notoriety during the Second World War, when civilians sheltering in the London Underground railway tunnels detailed being bitten [43]. Later, during the 1990s, *Cx*. *molestus* were reported once again in London Underground tunnels, as well as in other subterranean urban environments, such as sewage treatment facilities [17]. However, despite their historic presence, there have been no recent reports of mosquito biting on the London Underground (Representatives from Transport for London, personal communication, 2024). A more likely threat to public health could come from above-ground biting populations [44], which may increase in number in the case of urban flooding after heavy rainfall [45]. However, since mammals are dead-end hosts, for zoonotic transmission to occur, *Cx. molestus* would need to first bite birds involved in USUV transmission [32]. According to our study, this would also require sustained high average temperatures of around 23°C over several weeks, suggesting the public health threat of *Cx. molestus* in London is currently low.

On the other hand, hybridization events between *Cx. molestus* and *Cx. pipiens* could lead to populations with broader feeding habits, targeting both birds and humans [18–20,46]. Eleven hybrids were identified from both under- and over-ground catches in this study suggesting such admixture to some extent is already occurring. However, only five individuals became infected with none demonstrating USUV saliva presence suggesting further infection studies are required possibly by finding sites with larger hybrid numbers or by experimentally crossing the two biotypes.

The patterns of *Cx. pipiens* infection and transmission for this study are less clear compared to *Cx. molestus*. Although infection rates for all four temperatures assessed generally remain low to moderate, infection rates at 14 dpi appear significantly different at 19°C and 23°C compared to 17°C and 21°C with transmission potential also observed in the former interventions. The lack of a clear effect of temperature is likely due to a few methodological differences which separate the *Cx. pipiens* and *Cx. molestus* experiments. First, *Cx. molestus* were collected at a single time point from a single water body. In contrast, due to poor feeding rates of field-caught *Cx. pipiens*, multiple catches were pooled over several months from various water bodies at two sites in northern England in an attempt to get matched data comparable to *Cx. molestus*. Second, in attempts to mimic natural environmental conditions, immatures were kept in an outdoor insectary where temperatures varied with the ambient outdoor temperatures compared to *Cx. molestus* which were kept in a standardised climate-controlled indoor insectary. This introduces possible variation between populations through developmental [47,48], genetic [49–51] and microbiome effects [52–54], which could explain the variability in infection and transmission rates of *Cx. pipiens* observed across different temperatures. To overcome this, the authors suggest using a blood-soaked cotton feeding method [30] in future studies to increase feeding rates and reduce the impacts of these environmental and genetic biases by relying on fewer catches.

Understanding the thermal limits of USUV transmission is crucial for predicting and mitigating future outbreaks. The confirmation of USUV transmission by *Cx. pipiens* at temperatures as low as 19°C is notable. The widespread presence of this common (and potentially major vector) species in the UK suggests that USUV may impact animal health beyond the currently recognized endemic areas, with future rising temperatures potentially facilitating its spread [55,56]. Most previous USUV vector competence studies involving *Cx. pipiens* have been conducted at higher temperatures which are not comparable to the temperatures observed in cooler regions of the UK [26,29,31,57]. Therefore, despite USUV being currently restricted to regions in the south of England, our results suggest the potential viable spread of USUV to other regions within the thermal limits of the virus. Specifically, areas as far north as Yorkshire, and the North West of England could become new regions of concern, having potential implications for surveillance and resource allocation. For example, increased USUV testing of dead birds and mosquito populations in these areas could provide valuable information on the virus’ spread. Additionally, targeted recruitment for citizen science projects, such as the British Trust for Ornithology’s (BTO) survey of blackbirds in gardens [58], could provide an opportunity to enhance monitoring efforts by tracking bird declines in these areas that could potentially be linked to USUV.

Intriguingly, the only *Cx. pipiens* time point and temperature to have a 0% infection rate was at the latest time point (28 dpi) and warmest temperature (23°C). It is possible that this observation of a decreasing infection rate could be due to mosquitoes clearing the virus at later time points and higher temperatures. This has been suggested by Chapman *et al.* [59] where field-caught mosquitoes used from the same area as this study and at a similar temperature (24°C) often led to reduced transmission rates and titres for Japanese encephalitis virus (JEV) and Ross River Virus later in infection. Alternatively, infected mosquitoes under these conditions could be more likely to die and be excluded from analysis (i.e., survival bias). When considered as a whole, 4.6% of the total *Cx. pipiens* assessed across all temperatures (*N* = 237) produced expectorate with viral RNA. This is slightly higher than other studies using field-caught *Cx. pipiens* which found low transmission rates between 0 and 1.4% [28,29,60]. The low transmission rates observed in this study and elsewhere suggest other species should be monitored for their role in outbreaks. Additional species to interrogate include *Culex modestus* which has been shown to be competent for USUV in Belgium [28], where in the same study *Cx. pipiens* lacked transmission potential. *Cx. modestus* is currently found in the South East and East of England, occupying estuarine and coastal areas of Kent, Essex and Suffolk [61]. Therefore, in regions where both species coexist, this may exaggerate the impact of *Cx. pipiens* on USUV transmission dynamics [62].

Another species of interest is *Culiseta annulata*, which has a similar ecological niche to *Culex* spp. and has been known to feed on birds [63], larger mammals [64,65] and humans [63,66]. To date, the only implication of vector potential for *Cs. annulata* is from the detection of USUV in collected pools from Italy and Austria [67,68]. Our observation that *Cs. annulata* exhibits competence for USUV is the first to suggest their potential involvement in transmission cycles. The detection of elevated virus titres in the saliva of a single individual compared to *Cx. molestus* and *Cx. pipiens* could indicate an increased likelihood of USUV transmission during a single biting event.

Similarly, *Aedes detritus* has previously been implicated as a vector transmitting Japanese encephalitis virus (JEV) and WNV [59,69], and are of concern for public health as a result of their voracious human biting habits [32,70]. A single pool of *Ae. detritus* was previously found to contain USUV in the Molise region of Italy [67]. However, the lack of infection at UK temperatures in indigenous *Ae. detritus* suggest this population is unlikely to currently pose a risk to animal or public health. Further competence studies at higher temperatures are required to assess the vector potential of *Ae. detritus*’ for future UK climate scenarios.

In conclusion, this study is the first to report on the transmission potential of field-caught mosquitoes under current climatic conditions, using a circulating strain of USUV relevant to the UK context. We describe *Cx. molestus* as a competent vector that should be monitored for future public health threats in London, although the current risk appears low due to their need to first access infected birds and the requirement for sustained high temperatures for USUV transmission. Additionally, we identify UK regions within the thermal limits of the virus where *Cx. pipiens* populations have the potential to transmit USUV. Finally, we report the first evidence that *Cs. annulata* is able to transmit USUV.

## Author contributions

MSCB and MB contributed to the conceptual development of the project and acquired funding. JP, EW, AV, KS, and JM carried out fieldwork and collected samples. KLM and NJ provided and titrated virus. JP, EW, KS and JTH assisted in rearing of mosquitoes. JP and NS carried out vector competence experiments. JP and JTH assisted in RNA, DNA extractions. SM generated temperature suitability maps. JP analysed the data and led the writing of the manuscript with assistance from all authors.

## Acknowledgements

The authors would like to thank Dr Peter Enevoldson for providing *Aedes detritus*.

---

**Supplementary Figure S1.**
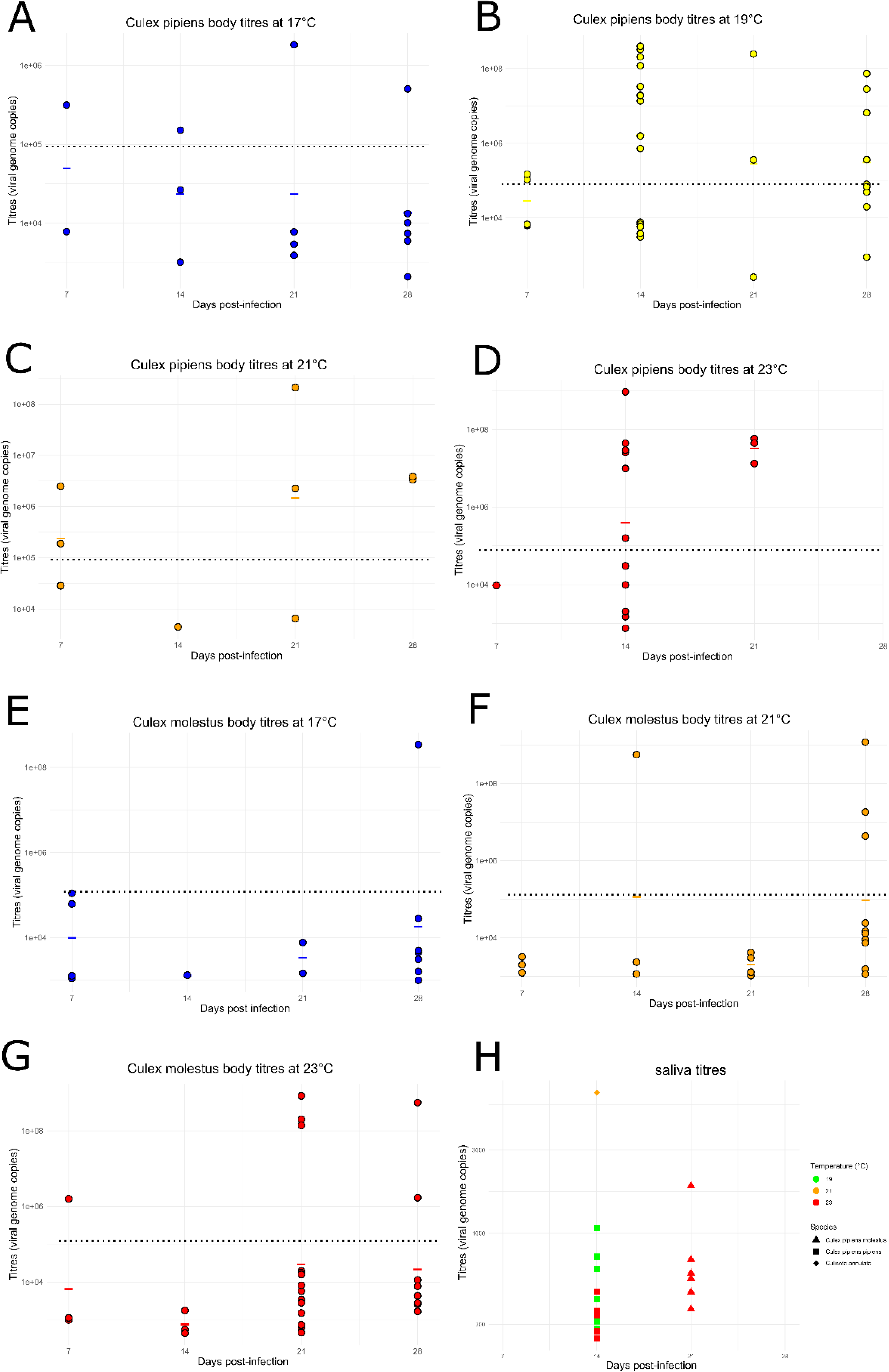
Usutu virus body titres at temperatures 17°C, 19°C, 21°C, 23°C for *Cx. pipiens* (A-D) and *Cx. molestus* (E-G) with saliva titres for each species and temperature (H). Horizontal coloured lines indicate titre means and dotted lines indicate mean titre of USUV-infected mosquito controls at 0dpi.

