## Supplementary figures and images for "UK mosquitoes are competent to transmit Usutu virus at native temperatures"

### Supplementary Figure S1

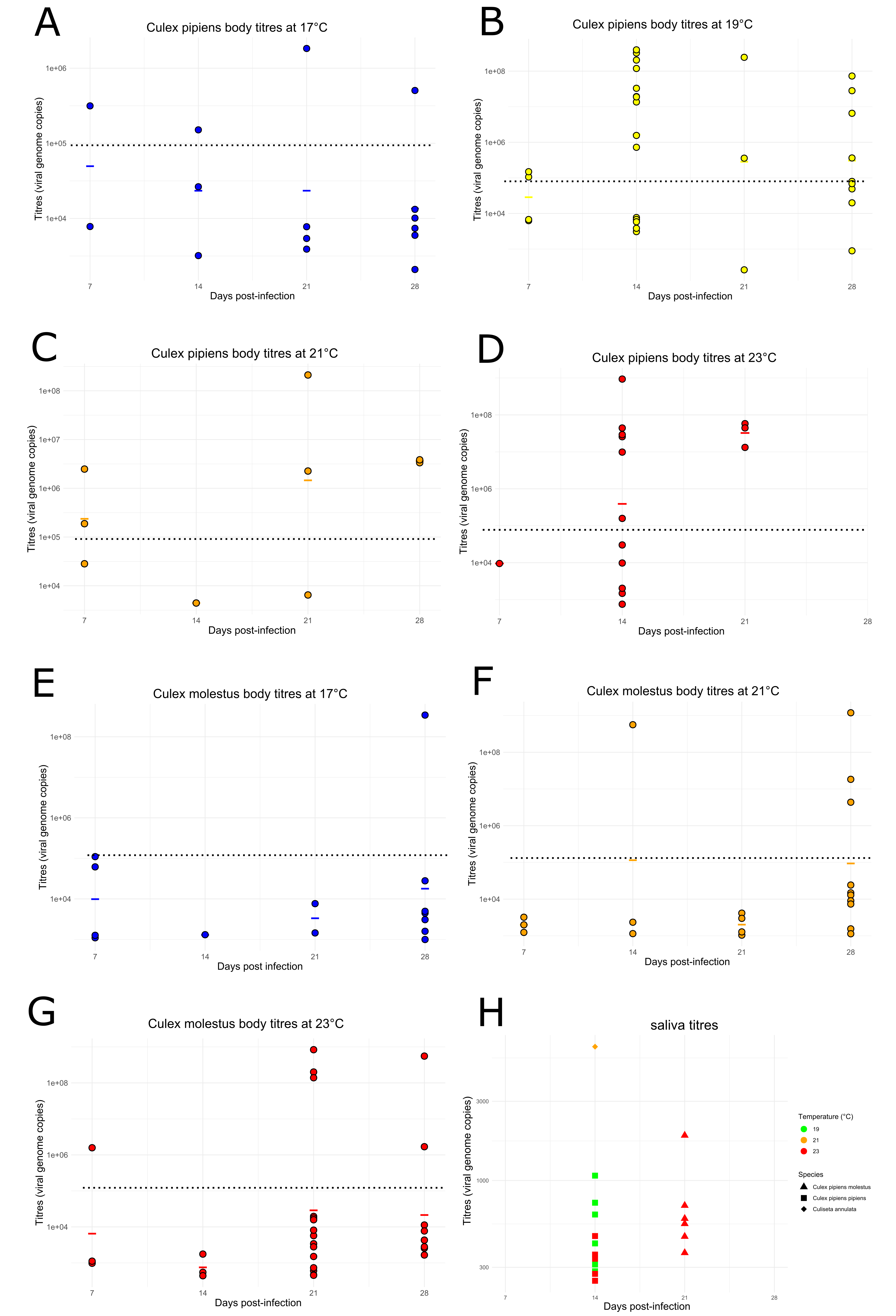
